# Sleep deprivation disrupts error awareness and subsequent behavioural regulation

**DOI:** 10.64898/2026.03.04.709320

**Authors:** Hans Kirschner, Jana Tegelbeckers, Daniel Janko, Luisa Göde, Markus Ullsperger

## Abstract

Sleep deprivation is known to impair cognitive performance, yet its effects on error awareness and subsequent behavioral adjustments remain incompletely understood. Here, we investigated how sleep loss affects the use of subjective performance evaluation to guide post-error adaptations.

Thirty healthy adults completed a novel, gamified error awareness multi-rule Simon task once while well rested and once after 24 h of total sleep deprivation. On each trial, participants reported both their task response and subjective evaluation of response accuracy. This design allowed us to dissociate objective performance from subjective error awareness and to examine their influence on subsequent behavior over time.

Sleep deprivation slowed responses, reduced accuracy, increased missed responses, and decreased the proportion of consciously detected errors. These effects increased with time on task and were accompanied by greater instability in sustained attention. Critically, post-error adjustments were driven by subjective error awareness rather than factual error commission. Reaction times slowed most strongly after subjectively perceived errors, including instances in which the preceding response had been objectively correct. Accuracy showed post-error decreases that were most pronounced following unaware errors. Sleep deprivation further altered these awareness-dependent control processes, particularly in later task phases.

Together, these findings indicate that sleep deprivation disrupts both error awareness and the effective use of awareness signals for behavioral regulation.

**Statement of significance:** One night of total sleep deprivation reduces behavioral error awareness and disrupts post-error adjustments in a time-dependent manner. Crucially, our findings show that adaptive cognitive control is strongly shaped by subjective error awareness—even when that awareness is inaccurate. By identifying conscious performance evaluation as a key mechanism linking sustained attention, sleep loss, and behavioral regulation, this work highlights the importance of considering subjective awareness when studying adaptive control under fatigue.

## Intro

Loss of sleep has detrimental effects on cognitive performance by impairing alertness, slowing down responses and rendering mistakes more likely (Chee et al., 2008). The significance of these effects becomes evident in drowsiness-related car crashes (Anderson et al., 2023) and fatigue-related errors in shift working medical personnel (Lockley et al., 2007). In these cases, the monitoring of behaviour, detection of mistakes and subsequent behavioural adaptation is of utmost importance. Yet, relatively little is known about the effects of sleep deprivation on the ability to consciously detect errors (typically called error awareness).

So far, the scientific literature presents an ambiguous picture of the relationship between insufficient sleep and performance monitoring (as reviewed by Boardman et al. (2021)). Some studies indicate reduced monitoring accuracy and increased response times (Anderson & Platten, 2011; Cain et al., 2011; Drummond et al., 2006; Hsieh et al., 2007; Hsieh et al., 2010; Tsai et al., 2005b) while others find no differences between sleep deprived and well rested performance on self-evaluation (Asaoka et al., 2010; Murphy et al., 2006; Schapkin et al., 2006).

Error detection is a central component of performance monitoring and crucial for goal-directed behaviour. Electrophysiological components are commonly used to measure brain responses to errors. After sleep deprivation, the error-related negativity (ERN) and the error positivity (Pe) have been shown to be reduced (Boardman et al., 2021), indicating both preconscious (ERN) and aware (Pe) error detection to be diminished. However, these studies fail to relate these brain responses to behavioural awareness measures. To bridge this gap, tasks are needed which measure individual performance monitoring and cause sufficient numbers of aware and unaware errors (Fischer et al., 2017). Only one study so far has used such a task in the context of sleep deprivation: Using the Error Awareness Task (EAT, Hester et al., 2005), Boardman et al. (2024) reported no effect of 24 h of total sleep deprivation on either behavioural or electrophysiological measures of error awareness.

The original EAT is a complex task during which participants have to monitor multiple task rules regarding different Go/NoGo conditions. It measures response inhibition errors which have to be detected by pressing an “error button”. This procedure might lead to particular task effects because 1. errors of commission cannot be corrected, 2. trials for which the error button was pressed differ from all other trials, and 3. participants do not have to respond on every trial (Kirschner et al., 2021). Therefore, the EAT is not optimal to investigate error awareness-related changes in post-error adaptation of behaviour.

After error commission, various post-error adjustments occur (Danielmeier & Ullsperger, 2011). These adjustments can manifest as increased reaction times after an error (post-error slowing, PES) or as improved accuracy on subsequent trials (post-error increase in accuracy, PIA). Both phenomena are thought to reflect general performance-adjustment mechanisms, as they have been observed across a wide range of tasks (Cavanagh et al., 2009; Gehring & Fencsik, 2001; Nieuwenhuis et al., 2002). However, while many studies have reported performance improvements following errors (Fischer et al., 2015; Grützmann et al., 2014; Seifert et al., 2011), other work has documented performance declines (Adkins et al., 2024; Houtman & Notebaert, 2013).

A long-standing question has been whether these post-error adjustments depend on error awareness (Kirschner et al., 2021). Evidence in the literature is mixed. In a well-rested state, post-error slowing is not always found which may result from long inter-trial intervals (Danielmeier & Ullsperger, 2011) and small sample sizes. But when it was found, it seemed that post-error slowing only occurs after aware errors (Cohen et al., 2009; Endrass et al., 2012; Kirschner et al., 2021; Klein et al., 2007; Nieuwenhuis et al., 2001; Wessel et al., 2011). However, not all studies demonstrate this relationship (Endrass et al., 2007; Wessel et al., 2011 Exp.1). The association between post-error improvement in accuracy and error awareness is rarely reported, and when it is, it appears only after aware errors (Klein et al., 2007). Notably, post-error adjustments have been shown to be impaired by sleep deprivation (Tsai et al., 2005a; Y. Zhang et al., 2025) but to our knowledge no study to date has investigated how error awareness might modulate this relationship.

We therefore aimed to examine how one night of total sleep deprivation affects behavioural error awareness and subsequent adjustments in task behaviour. We developed a novel gamified Simon task that allowed us to test for these effects within one experiment. We found that total sleep deprivation diminished task performance and reduced error awareness. But most importantly post error adaptations depended on perceived task performance and were thus also altered by sleep deprivation. Adaptive control therefore seems to rely on awareness signals which in turn require attentional resources. This is further supported by an increase in sleep deprivation effects with the time spent on task.

## Methods

### Participants

Thirty-five participants took part in an initial screening session, during which they received detailed information about the study’s purpose and procedures and provided written informed consent after eligibility was confirmed. Eligibility criteria included: age between 18 and 40 years, no current or past neurological or psychiatric disorders, eligibility for fMRI, fluent German language skills, normal or corrected-to-normal vision, and regular sleep patterns (i.e., no night-shift work).

Three participants performed below chance level in at least one session and were excluded from further analyses. Two additional participants did not complete the Error Awareness Task due to technical issues. The final sample therefore consisted of 30 participants (13 male, 17 female), who each completed a novel Error Awareness Simon Task (EAST) once after total sleep deprivation and once while well-rested.

### Procedure

After the screening, participants were paired in groups of two and completed two study appointments during which they underwent different behavioural tests and an fMRI session (the fMRI data were not part of the current study), once in a wakeful rested state (WR) and once after 24h of total sleep deprivation (TSD). These sessions were one week apart and the order was counterbalanced. Half of the participants started with TSD, the other with WR. Before and between test sessions participant’s daily activity and sleep patterns were monitored by Fitbit Charge 5 devices, to ensure that the days leading up to the test sessions were comparable. These data showed that participants slept about 8h on average (WR: 7.9h ± 0.89, TSD: 7.85h ± 0.83), with similar Fitbit sleep scores (WR: 74.95 ± 4.35, TSD: 76.05 ± 4.64) and comparable amounts of daily steps per week (WR: 10367 ± 4934, TSD: 10960 ± 5693.9).

The WR session was scheduled for 8am with one participant starting in the fMRI scanner and the other starting with the error awareness task. Before these experiments, participants performed a modified version of the Psychomotor Vigilance Task (PVT; Dinges & Powell, 1985) and rated their subjective level of sleepiness on German translations of the Karolinska Sleepiness Scale (KSS; Shahid et al., 2012) and the Epworth Sleepiness Scale (ESS; Johns, 1991). The TSD session began at 10pm the previous night. The pair of participants and one supervisor spent the night at the Leibniz Institute for Neurobiology. Every 2h participants completed the PVT, KSS and ESS. Between these tests, they were free to entertain themselves with light activities, e.g. board games, puzzles, watching TV. At about 6am both participants performed the error awareness task.

Upon study completion, participants were reimbursed with a total of 150€. Our procedure is in line with the Declaration of Helsinki and was positively evaluated by the local ethics committee of the Otto von Guericke University Magdeburg.

### Tasks

#### Error Awareness Simon Task (EAST)

We employed a novel multi-rule Simon task to measure error awareness and cognitive control processes (see Figure 1). On each trial, participants were instructed to respond as quickly and accurately as possible according to the colour (blue or green) of a circle presented either on the right or left side of the screen (blue = inner left response button, green = inner right response button; colour-response-mapping counterbalanced across subjects). Trials were congruent when stimulus and requested response were on the same side and incongruent when they were on opposite sides. To induce a sufficient number of errors, we introduced two additional one-back rules. Here, participants had to track the identity of the colour circle from the last trial. First, if there was a direct colour repetition, participants had to press the outer left or right button, according to the current trial’s circle colour. Second, if a dark colour follows a bright colour, participants also had to press the outer left or right button, according to the current trials circle colour.

**Figure 1.**
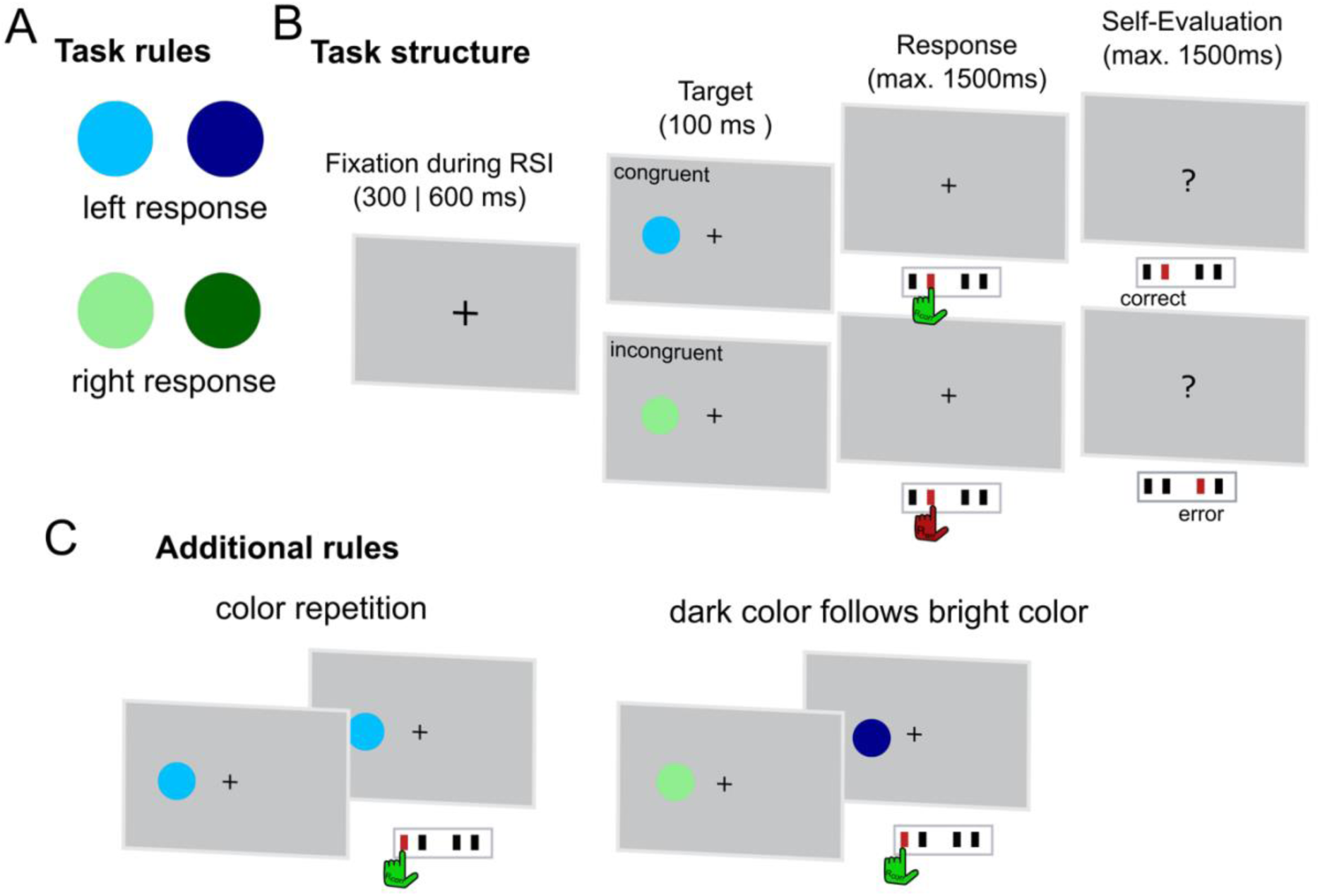
A novel multi-rule reaction time task to measure error awareness and cognitive control processes. **(A)** Basic rules of the task. In each trial participants have to press the left button for blue colours and the right button for green colours. **(B)** Time course of the task. Each trial began with the presentation of a fixation cross (Response Stimulus Interval (RSI)) for either 300ms or 600ms. Hereafter, a coloured circle was presented either on the left or right side of the fixation cross. After the target disappeared, participants had to respond to the colour of the circle within a response window of max. 1500ms. If participants failed to respond in time, they were instructed to speed up and the next trial started. If participants responded, they were presented a question mark, indicating that they had to evaluate the accuracy of the previous response. **(C)** Additional task rules. To induce a sufficient number of errors, we introduced additional rules to the task. Here, participants had to track the identity of the colour circle form the last trial. If the colour was repeated, participants had to press the outer left or right button, according to the current trials circle colour (left illustration). If the previous trial displayed a bright colour and the current trial a dark colour, participants also had to press the outer left or right button, according to the current trials circle colour (right illustration). These special rule trials comprised 40% of the trials in the task (20% colour repetition trials (n=200); 20% dark colour follows bright colour trials (n=200)).

After each response (i.e., the primary task), participants had to indicate whether they thought they responded correctly or not (i.e., secondary task) by pressing either the left button (subjective correct response) or right button (subjective error). The self-evaluation response-mapping was counterbalanced across subjects.

Each trial began with the presentation of a fixation cross (Response Stimulus Interval (RSI)) for either 300ms or 600ms. Hereafter, a coloured circle was presented one the left or right side of the fixation cross. After the target disappeared, participants had to respond to the colour of the circle within a response window of max. 1500ms. If participants failed to respond in time, they were instructed to speed up and the next trial started. If participants responded, they were presented a question mark, indicating that they had to make their accuracy judgment (max 1500ms). In total, the task comprised 1000 trials (50% incongruent). Secondary rule trials comprised 40% of the trials (20% colour repetition trials (n=200); 20% dark colour follows bright colour trials (n=200)). Trials were presented in 10 blocks of 100 trials, with a short self-paced break in between blocks. To ensure task motivation, in each break, participants were presented their current task score along with their personal high score. The task score was calculated according to the following formula:

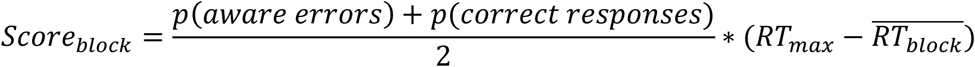

where *p*(*aware erros*) is the proportion of aware errors among all errors in a given block, *p*(*correct responses*) is the proportion of correct responses among all trials in a given block, *RT*_*max*_ is the response window of 1500ms, and 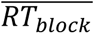 is the mean RT in a given block. Participants were informed how the score was calculated before the task.

In each block, participants were instructed to try to beat their personal high score. Before starting the experiment, participants performed an extensive training to ensure task understanding.

#### PVT

To measure sustained attention, we used a modified version of the Psychomotor Vigilance Task (PVT; Dinges & Powell, 1985). Participants were asked to press the space button as fast as possible when a square appeared in the centre of the screen. In most cases, and for the first ten trials this square was always red. Afterwards, 15 oddball targets in different colours were randomly intermixed. The task took 10min and the time between targets ranged between two and ten seconds.

### Questionnaires

The Karolinska Sleepiness Scale (KSS) assesses the individual level of sleepiness on a nine-point scale, ranging from 1 = “extremely alert” to 9 = “very sleepy, great effort to keep awake”. Similarly, the Epworth Sleepiness Scale (ESS) measures “daytime sleepiness” by asking people to rate their perceived likelihood to fall asleep while they were engaged in eight different activities. A 4-point rating scale is used ranging from 0 = “would never nod off” to 4 = “High chance of nodding off”. For both scales higher values indicate higher degrees of sleepiness.

### Data analyses

#### Effects of sleep deprivation on task performance

We first examined the effects of sleep deprivation on overall task performance indices (i.e., accuracy, error awareness, missed responses, and reaction time) by fitting a set of Bayesian linear mixed-effects models. For each subject s, the model predicted the respective task performance index:

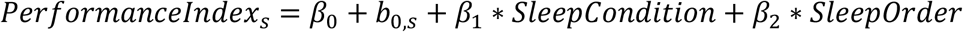

where *βs* are fixed effects, *b*_s_ denotes the subject-specific random intercepts, SleepCondition indicates the state in which the task was performed (coded as −1 for well-rested and 1 for sleep-deprived), and SleepOrder is a control regressor accounting for session order (sleep deprivation in the first vs. second session, coded as −1 and 1, respectively).

Using the same approach, we next identified the critical factors influencing reaction time (RT) and accuracy in more detail. We added task congruency (congruent vs. incongruent trials, coded −1 vs. 1), accuracy (correct vs. error, coded −1 vs. 1, only RT model), subjective (perceived) accuracy (SubAccuracy; correct vs. error, coded −1 vs. 1), trial type (standard vs. secondary rule, coded −1 vs. 1), response stimulus interval (RSI; short vs. long, coded −1 vs. 1) and trial number (reflecting time on task effects) to the existing model. Moreover, post-error adaptations were included by assessing the effects of previous accuracy (*Accuracy*_*t*−1_) and subjective previous accuracy (*SubAccuracy*_*t*−1_).

The regression models on RT:

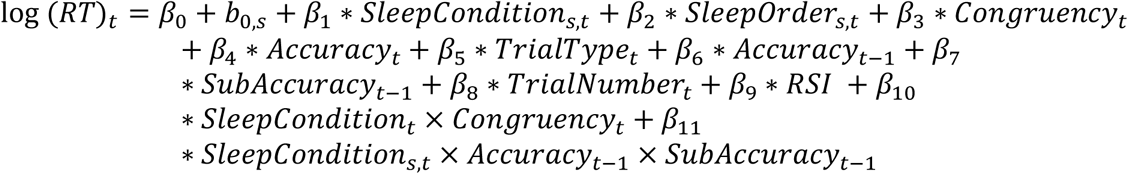

The logistic regression models on accuracy:

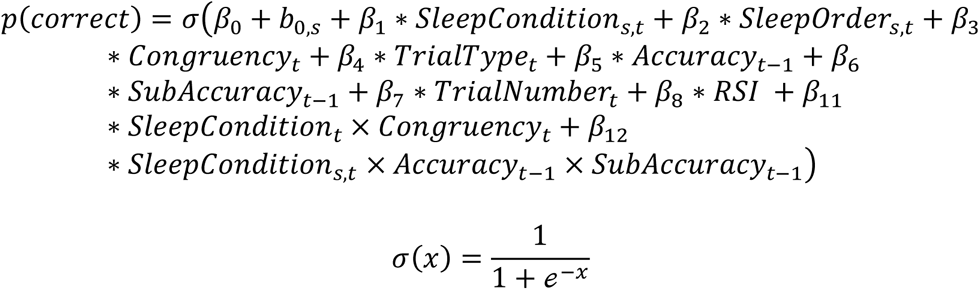

For all regression models, we report fixed effects as the mean of each parameter’s marginal posterior distribution together with 95% credible intervals, representing the range containing 95% of the posterior density. Parameter values outside this range are improbable given the model, data, and priors. When the credible interval excludes zero, we interpret this as evidence for a meaningful effect.

##### Self-evaluation performance analyses

We assessed self-evaluation performance within the signal detection framework (Green & Swets, 1966; Swets, 2014). Specifically, we estimated the signal detection theory parameters stimulus sensitivity (d′) and response criterion (c) to quantify participants’ ability to evaluate their own responses and any potential bias in this evaluation. These indices were derived from the observed hit and false alarm rates of the self-evaluation responses as follows:

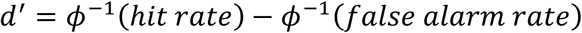

and

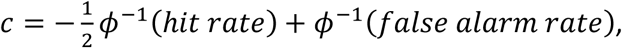

Here, *ϕ*^−1^is the inverse of the cumulative standard normal distribution that transforms hit rate and false-alarm rate into z scores. Hit rate and false-alarm rate were defined by treating the errors in the primary task as target. Therefore, negative c values indicate a bias to under report errors, whereas positive c values indicate a bias to over report errors. We then examined the receiver operating characteristic (ROC; Luce, 1963) curves (see Figure 3).

#### Time-on-task effects analyses

To examine how sleep deprivation affected task performance over the course of the task, we first computed the variance time course (VTC) of response times: we z-transformed RTs within each participant to normalize the values. We then calculated the absolute value of the z-scores, where higher values indicated greater deviations from the mean (including both very slow and very fast responses), while lower values reflected RTs closer to the task mean. Next, we computed a smoothed density estimate on the z-scored RT data using a Gaussian kernel with a full width at half-maximum (FWHM) of 9 trials, integrating information from the surrounding 20 trials through a weighted sum. Although the VTC provides a continuous within-subject measure of variability, segmenting the time series into high- and low-variability trials allows for direct accuracy comparisons between more and less stable periods of performance. To classify trials, we performed a median split on the smoothed VTC, defining low-variability trials as “in the zone” and high-variability trials as “out of zone,” based on prior research (Esterman et al., 2012).

Next, we computed smoothed density estimates of key task performance variables (error rate, error awareness rate, and VTC) using a Gaussian kernel with a full width at half maximum (FWHM) of 9 trials. This procedure effectively integrated information from the surrounding 20 trials via a weighted average of the performance measures. Then, we discretized the available trials into three quantiles, estimated the area under the ROC curve (AUC) using a trapezoidal approximation (trapz function in MATLAB) and plotted the difference (Δ = WR − TSD) to visualize these time-on-task effects (see supplementary results for a more fine-grained discretization using a sliding window approach).

#### Posterior inference and model fitting

All regression models were fit using No-U-Turn Sampling (Hoffman & Gelman, 2014) as implemented in the BAMBI toolbox (Capretto et al., 2022) in Python. Four chains with 3000 samples (1000 discarded as burn-in) were run for a total of 8000 posterior samples per model. Chain convergence was determined by ensuring that the Gelman–Rubin statistic *R* was close to 1 and by visual inspection of trace plots. Default weakly informative priors implemented in the BAMBI package (Capretto et al., 2022) were used for each regression model.

### Data availability

All code and behavioural data in this manuscript along with the error awareness task can be downloaded on the Open Science Framework at [INSERT LINK AFTER PUBLICATION].

## Results

### Effects of sleep deprivation on subjective sleepiness levels and sustained attention Sleep deprivation enhanced sleepiness and diminished vigilance

Total sleep deprivation (TSD) increased the subjective level of sleepiness compared to the well-rested (WR) condition before the error awareness task (Figure 2, KSS: *β*_*SleepCondition*_ = 1.51, 95% CI = [1.21, 1.82]; ESS: *β*_*SleepCondition*_ = 3.62, 95% CI = [2.83, 4.35]). While participants‘ reaction times and reaction time modulation to standard vs. oddball stimuli in the PVT were comparable (all 95% CIs included zero), they missed more responses after sleep deprivation (misses mean/SEM SD: 47.2/3.87 vs. 40.76/3.68); *β*_*SleepCondition*_ = 3.199, 95% CI = [0.61, 5.65]).

**Figure 2.**
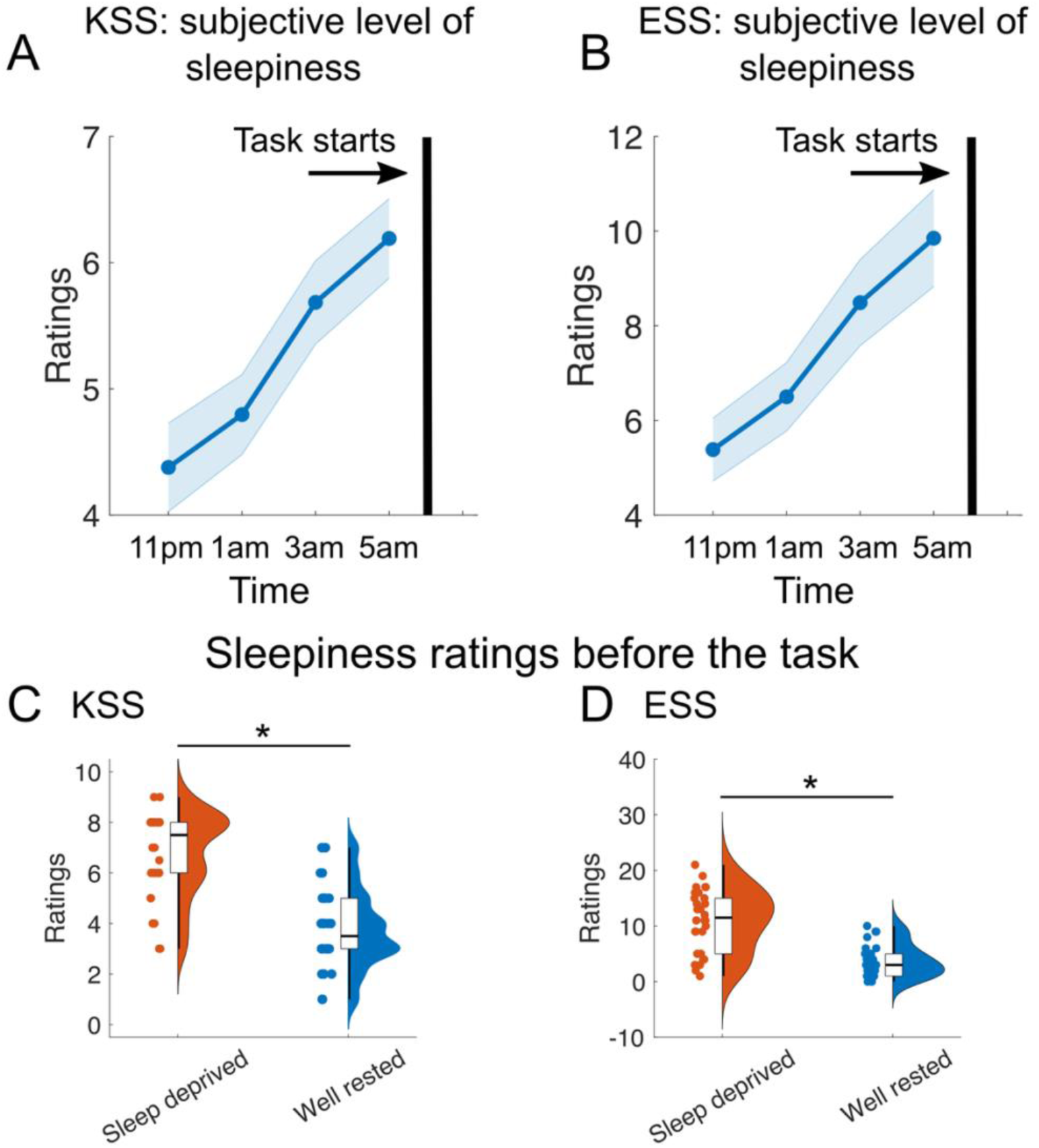
Sleep deprivation significantly increased the subjective level of sleepiness. Sleep deprivation significantly increased subjective level of sleepiness on German translations of the Karolinska Sleepiness Scale (KSS; panels **A/C**) and the Epworth Sleepiness Scale (ESS; panel **B/D**). The vertical black line indicates the time at which the task was completed. The asterisks indicate meaningful effect of TSD (i.e., credible interval excludes zero). Shadings in A and B indicate standard error of the mean.

### Sleep deprivation impairs overall task performance

Overall, participants performed the task with an error rate of ∼10%. We found that participants made more errors when they were sleep deprived (error rates in % (mean/STD): sleep deprived: 12.20/5.59 vs. well rested: 9.12/5.31; *β*_*SleepCondition*_ = 1.55, 95% CI = [0.73, 2.30]). Moreover, sleep deprivation induced a higher proportion of missed responses (%misses (mean/STD): sleep deprived: 2.37/3.24 vs. well rested: 0.61/0.96; *β*_*SleepCondition*_ = 0.89, 95% CI = [0.43, 1.36]). Additionally, we found slightly higher overall reaction times in the primary task after sleep deprivation (RT in ms (median/STD): sleep deprived: 631.29/124.63 vs. well rested: 597.91/121.81; *β*_*SleepCondition* (log(*RT*))_ = 0.01, 95% CI = [0.001, 0.02]). Sleep deprivation did not affect response times during self-evaluation (RT in ms (medium/STD): sleep deprived: 249.71/116.44 vs. well rested: 224.95/129.53; *β*_*SleepCondition* (log(*RT*))_ = 0.003, 95% CI = [−0.01, 0.01]).

### Sleep deprivation reduces self-evaluation accuracy, while not affecting responses biases

A good discrimination between correct and erroneous responses was apparent across both testing sessions (see results related to d-prime below and Supplementary Figure 1). It is noteworthy, that our task induced a good proportion of unaware errors (on average n = 50 (STD=20) trials), rendering the task useful to study the effect of error awareness on cognitive control processes and neural correlates of performance monitoring (Fischer et al., 2017).

Formal analyses of self-evaluation performance within the signal detection theory are shown in Figure 3 and revealed three key insights. First, overall participants displayed excellent signal to noise discriminability. Generally, any d-prime value above zero indicates that the participant can discriminate the signal over noise above chance, with scores higher than two considered as very good discrimination (Swets, 2014). In our sample the average d-prime score was above 3 (d-prime mean/SEM: 3.08/0.18). Second, when comparing the effect of sleep-deprivation on d-prime across all trials, we did not find a credible effect (d-prime mean/SEM SD: 2.36/0.35 vs. WR: 2.73/0.36; *β*_*SleepCondition*_ = −0.18, 95% CI = [−0.51, 0.15]). Yet, follow-up analyses revealed reduced error awareness after sleep deprivation (*p*(aware errors) mean/SEM SD: 0.45/0.03 vs. WR: 0.52/0.03; *β*_*SleepCondition*_ = −3.34, 95% CI = [−5.23, −1.374]; see Figure 3A inset). Third, we found a consistent negative response bias indicating a general preference to not report errors, which in our task was mainly driven by very low false alarm rates (see Figure 3B and Supplementary Figure 1). Response bias was not affected by sleep deprivation (*β*_*SleepCondition*_ = −0.003, 95% CI = [−0.17, 0.17]).

**Figure 3.**
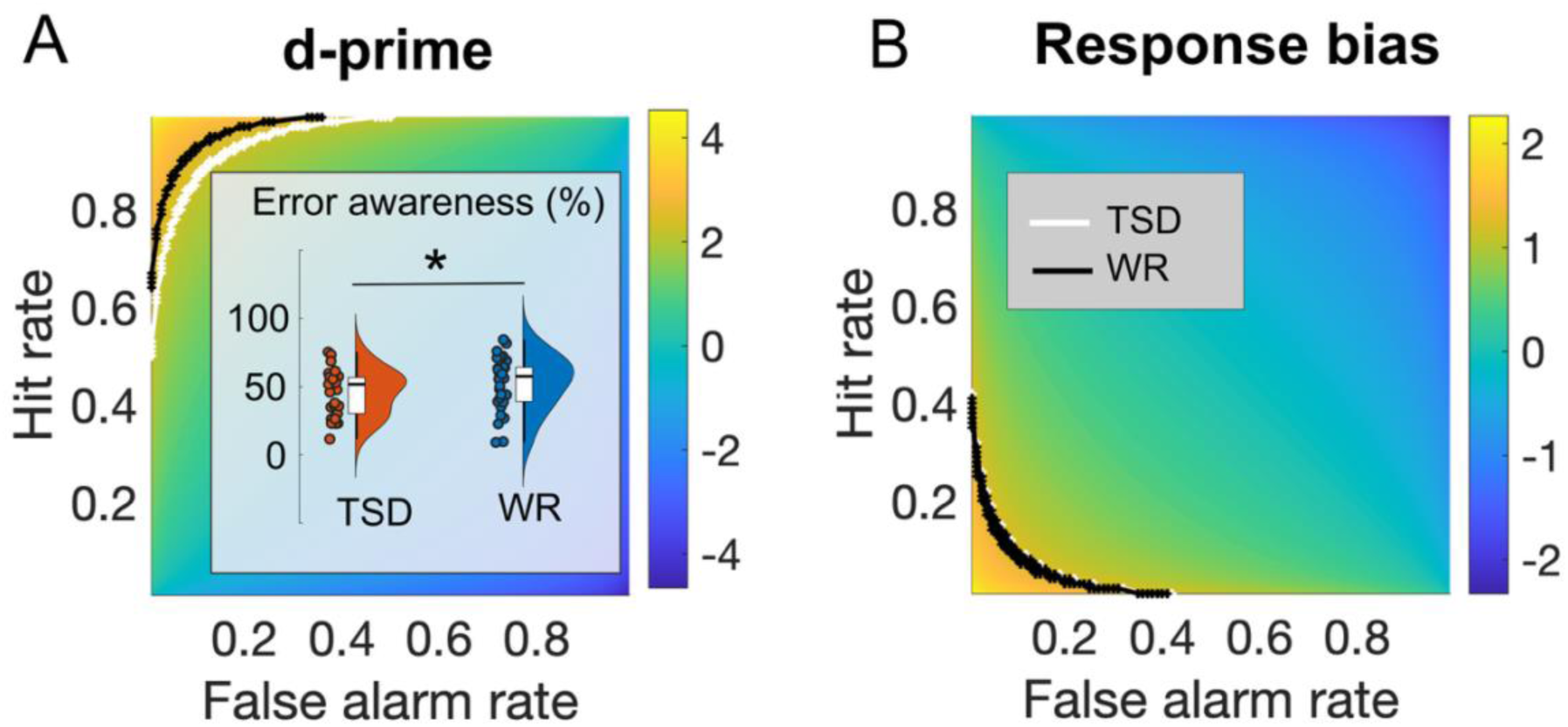
Sleep deprivation reduces self-evaluation accuracy, while not affecting response biases. Isosensitivity curves (ROC; Luce, 1963) for d-prime **(A)** and response bias **(B)** are drawn into the “yes” space of SDT. Specifically, we created a 2D space of p(Hits) and p(False Alarms) and drew the Isosensitivity curves for the average d-prime and response bias into this 2D space. This reveals how different combinations of p(Hits) and p(False Alarms) give rise to the same d-prime or response bias. Average scores for sleep deprivation are shown in white and scores for the well-rested condition are shown in black. D-prime values above zero indicate discrimination of signal over noise above chance, while scores higher than two are considered very good discrimination (Swets, 2014). Our results indicate that d-prime is descriptively reduced after sleep deprivation. Follow-up analyses revealed that this effect was mainly driven by a reduction in error awareness (see inset in **A**). The consistent negative response bias **(B)** indicates a general preference to not report errors, which in our task was mainly driven by very low false alarm rates. The asterisk indicates a credible effect of TSD (i.e., credible interval excludes zero).

### Sleep deprivation effects on performance and sustained attention depend on time-on-task

Next, we examined the time course of performance measures across the task by deriving smoothed density estimates. This yielded continuous trajectories of errors and unaware errors for each participant, capturing gradual changes in performance and error awareness over time (see Figure 4 A,B for individual examples).

**Figure 4.**
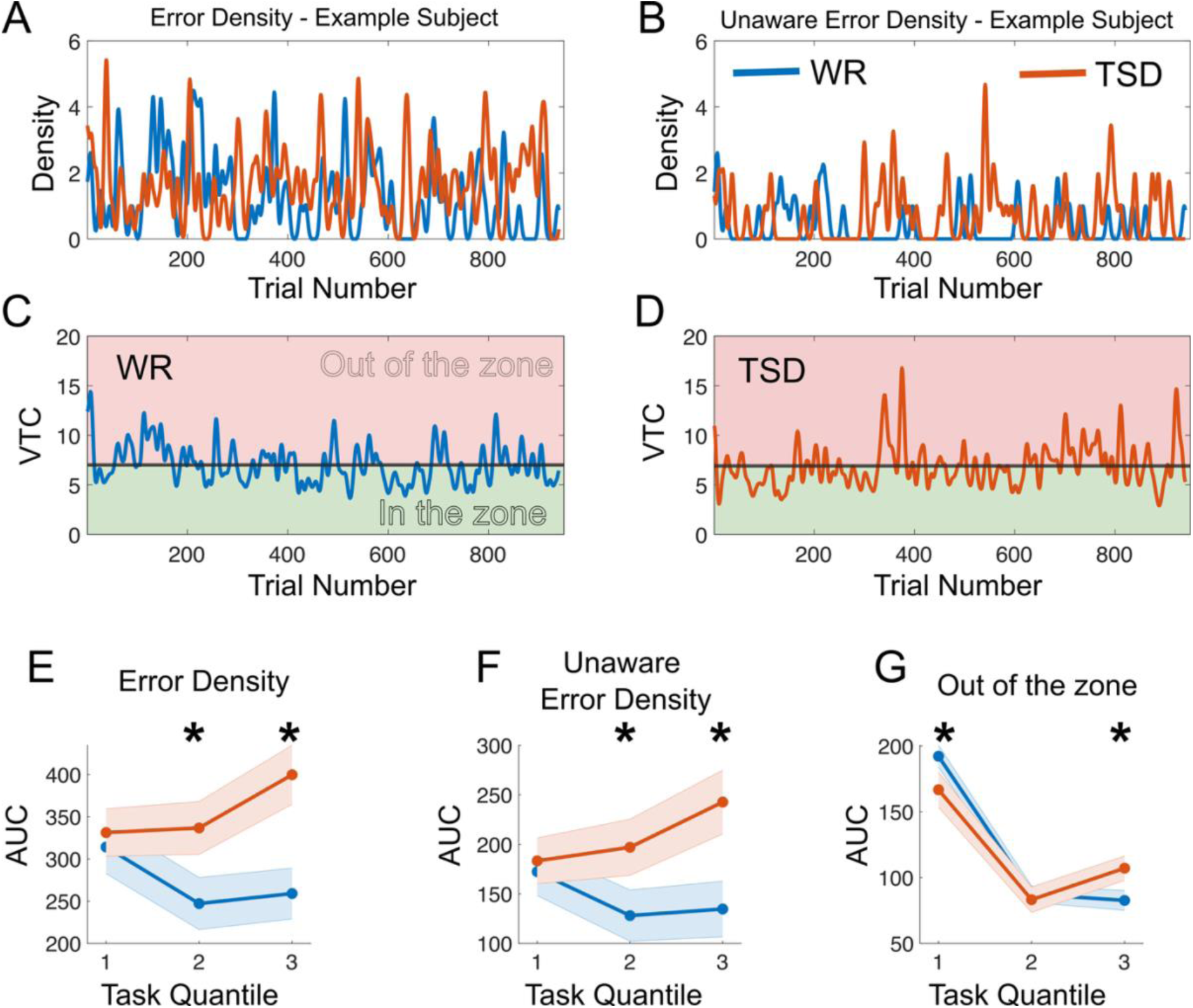
Sleep deprivation effects on performance and sustained attention depend on time-on-task. **(A-B)** Density measure trajectories for the number of errors **(A)** and unaware errors **(B)** for an example participant. **(C-D)** Variance time course (VTC) and unaware errors (vertical black lines) for two example participants: **(C)** well-rested and **(D)** total sleep deprivation. The VTC was calculated by averaging absolute deviations from the mean RT over a moving window of 20 trials. **(E-H)** Time-on-task effects on performance and sustained attention, assessed by dividing individual performance trajectories into three quantiles and estimating the area under the ROC curve (AUC) using a trapezoidal approximation (trapz function in MATLAB). The results indicate that total sleep deprivation primarily affects performance and attention in the later stages of the task. Note: WR = well-rested; TSD = total sleep deprivation; * = meaningful effect of TSD (i.e., credible interval excludes zero).

Building on prior work linking unaware errors to lapses in sustained attention (e.g., Harty et al., 2013), we quantified fluctuations in sustained attention over time using the variance time course (VTC; Esterman & Rothlein, 2019). The VTC provides a continuous, within-subject index of response-time variability, capturing alternations between periods of stable performance (“in the zone”) and unstable performance (“out of the zone”). Based on the smoothed VTC, trials were classified into low- and high-variability periods, allowing direct comparisons of accuracy between more and less stable attentional states.

To assess time-on-task effects on performance and sustained attention, we divided individual performance trajectories into three quantiles based on trial number. In the first quantile, error density and unaware error density were unaffected by total sleep deprivation (error density Q1: *β*_*SleepCondition*_ = −8.44, 95% CI = [−21.14, 37.71], unaware error density Q1: *β*_*SleepCondition*_ = 5.36, 95% CI = [−12.84, 23.64]). However, in the second and third quantiles, sleep deprivation credibly increased error and unaware error density (error density Q2: *β*_*SleepCondition*_ = 44.26, 95% CI = [19.51, 69.03], unaware error density Q2: *β*_*SleepCondition*_ = 34.25, 95% CI = [13.08, 54.03], error density Q3: *β*_*SleepCondition*_ = 70.33, 95% CI = [39.44, 99.15], unaware error density Q3: *β*_*SleepCondition*_ = 53.83, 95% CI = [32.98, 74.36]; see Figure 4E-F).

At the beginning of the task, well-rested participants spent more time in the “out-of-the-zone” attentional state than sleep-deprived participants (*β*_*SleepCondition*_ = −7.78, 95% CI = [−14.76, - 0.16]). Across the middle portion of the task, both groups showed a pronounced reduction in time spent out of the zone. In the final task quantile, the groups diverged, with sleep-deprived participants showing an increase in time spent out of the zone, whereas well-rested participants remained at a low level (*β*_*SleepCondition*_ = 7.15, 95% CI = [1.64, 12.83]; Figure 4G).

See the Supplementary Results for a more fine-grained discretization of time on task effects using a sliding-window approach.

### Post-error adjustments are affected by error awareness and sleep deprivation

To better understand how different task factors influenced participants’ behaviour—and to assess how post-error adjustments are affected by error awareness and sleep deprivation— we modelled the effects of key variables on reaction time and accuracy using two hierarchical regression models. Each model was applied to participants’ single-trial RT and accuracy data. The regression models were fitted hierarchically using Bayesian inference.

This analysis revealed several key insights, which are depicted in Figure 5. First, and in line with the overall performance analyses, sleep deprivation induced reliable slowing of responses (*β*_*SleepCondition*_ = 0.01, 95% CI = [0.006, 0.015]) and reduced accuracy (*β*_*SleepCondition*_ = −0.11, 95% CI = [−0.15, −0.08]). Second, we observed a congruency effect, with longer RTs (*β*_*Congruency*_ = 0.02, 95% CI = [0.017, 0.022]) and lower accuracy (*β*_*congruency*_ = −0.07, 95% CI = [−0.09, −0.04]) on incongruent trials compared to congruent trials. Sleep deprivation did not affect this congruency effect (i.e., 95% CI includes zero; see Figure 5A and C). Third, trial type (standard vs. secondary rule) did not affect RT (*β*_*TrialType*_ = 0.001, 95% CI = [−0.002, 0.003]), but did reduce accuracy (*β*_*TrialType*_ = −0.16, 95% CI = [−0.18, −0.14]). While sleep deprivation did not influence the effect of trial type on accuracy (*β*_*TrialType*:*SleepCondition*_ = −0.001, 95% CI = [−0.023, 0.024]), participants showed longer RTs on secondary-rule as compared to standard trials only when well rested (*β*_*TrialType*:*SleepCondition*_ = −0.004, 95% CI = [−0.006, −0.001]; see Figure 5B).

**Figure 5.**
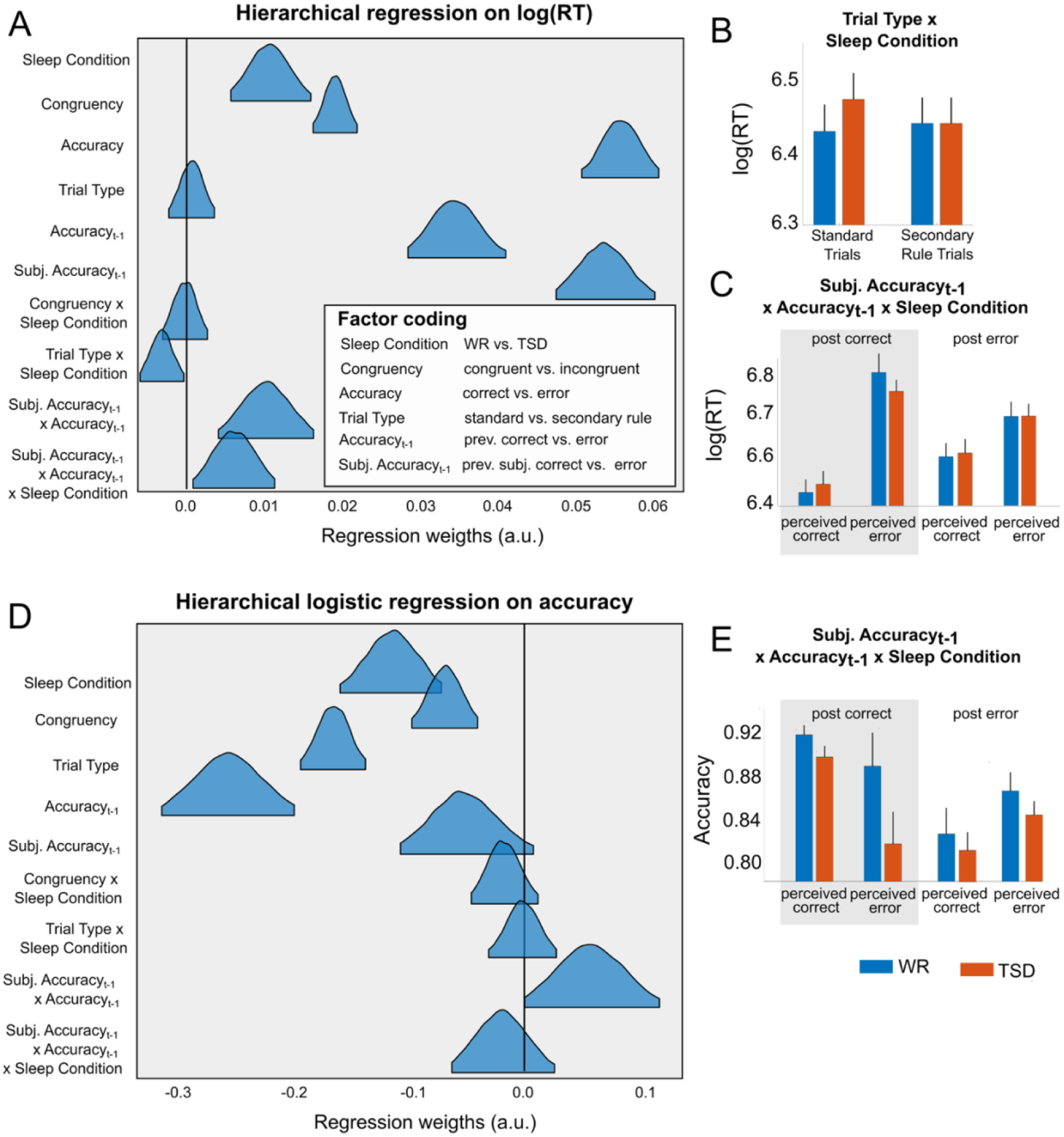
Sleep deprivation effects on performance and sustained attention depend on time-on-task. Panels **A** and **D** show posterior distributions of predictors from the hierarchical models on log-transformed response time (logRT) and accuracy. Panels **B**, **C**, and **E** display raw data splits to illustrate interaction effects. RT values reflect within-participant median RTs per condition (**B & C**).; accuracy values reflect within-participant mean accuracy per condition (**E**). Results demonstrate that sleep deprivation caused overall slower responses and reduced accuracy. A congruency effect was present—participants were slower and less accurate on incongruent trials—and this pattern was unchanged by sleep deprivation. Trial type affected accuracy, and it influenced RTs only when participants were well rested **(B)**. We also observed post-error adjustments: RTs increased after mistakes, and this slowing was even stronger when participants believed they had made an error, including in cases where their response had actually been correct (**C**). Accuracy showed post-error decreases, especially after unaware errors. Descriptively, after sleep deprivation, mistakenly believing one had made an error further impaired subsequent accuracy (**E**), although this effect was not statistically robust.

Furthermore, in the regression on the reaction time we found a triple interaction of sleep deprivation, previous accuracy, and previous subjectively perceived accuracy (*β*_PrevAcc:PrevSubjAcc:SleepCondition_ = 0.006, 95% CI = [0.002, 0.011]; see Figure 5A and C). This reflected two interesting findings: Reaction times were generally longer after errors (i.e., post-error slowing). But subjective awareness of having made a mistake had an even stronger effect. This was particularly true for correct responses that were misclassified as errors. In other words, participants slowed down most, when they believed that they had made an error, even and particularly when this belief was wrong. This effect was strongest in the WR condition. We also found post-error adjustments in the regression on accuracy. Specifically, we found no post-error improvement, rather a post-error decrease in accuracy, which was strongest for unaware errors (*β*_PrevAcc:PrevSubjAcc_ = 0.055, 95% CI = [0.005, 0.10]; see Figure 5D and E). Moreover, descriptively, after sleep deprivation subjectively perceived errors on actually correct responses were detrimental for subsequent accuracy (see Figure 5E). Yet, the 95% credible interval of the triple interaction of sleep deprivation, previous accuracy, and previous subjectively perceived accuracy included zero (see Figure 5D).

## Discussion

The present study examined how total sleep deprivation affects the awareness of committing errors and influences post-error behavioural adjustments. Using a novel, gamified multi-rule error awareness Simon task, we dissociated objective task performance from subjective error evaluation and traced their influence on subsequent behaviour over time. Three key findings emerged. First, sleep deprivation reduced the proportion of consciously detected errors. Second, these impairments increased with time on task and covaried with fluctuations in sustained attention. Third, we found that post-error adjustments were primarily driven by subjective error awareness rather than objective error commission, with perceived errors eliciting the strongest behavioural changes—even when the preceding response had been correct.

### Sleep deprivation, error awareness, and sustained attention

Consistent with prior work, 24h of total sleep deprivation led to comprehensive decline in overall task performance characterized by slowed responses, reduced accuracy, and an increased rate of missed responses, alongside elevated subjective sleepiness. While global self-evaluation sensitivity (d′) was not credibly reduced, explicit awareness of errors was significantly diminished. This dissociation suggests that sleep loss selectively reduces the likelihood that error signals cross the conscious awareness threshold rather than uniformly reducing metacognitive discrimination. Such a pattern aligns with electrophysiological findings showing attenuated error-related brain responses after sleep deprivation, particularly in later components associated with conscious error processing (Boardman et al., 2024; Boardman et al., 2021; Yue Zhang et al., 2025).

Critically, the effects of sleep deprivation were not static but increased with time on task. Using smoothed density estimates and the variance time course (VTC) as a behavioural measure of fluctuations in sustained attention (Esterman et al., 2012), we found that sleep deprivation primarily affected later task phases, with disproportional increases in error density, unaware error density, and time spent in an “out-of-the-zone” attentional state. These results converge with recent evidence that sleep loss amplifies attentional instability over time rather than producing a uniform performance decrement (Yang et al., 2025). One possible explanation for these behavioral fluctuations lies in an increased instability of the arousal system (Yang et al., 2025) but also the occurrence of “local sleep” episodes. These brief, region-specific neuronal off-periods occur in use-dependent cortical areas despite global wakefulness (Andrillon et al., 2019) and could contribute to attentional instability.

Together, our results support models of sustained attention that emphasize dynamic fluctuations between stable and unstable attentional states, which become more pronounced under prolonged wakefulness (Humphrey et al., 2026).

### The illusion of having made an error boosts post-error slowing

Beyond the detection of errors, adaptive control should use these signals to improve goal-directed behavior. Importantly, the present study demonstrates that these post-error adjustments depend more strongly on subjective belief of error detection than on actual error commission. We found that responses slowed following errors, consistent with classic post-error slowing (Danielmeier & Ullsperger, 2011) but crucially this slowing was even stronger following subjectively perceived errors, including cases in which the preceding response had been factually correct. The largest slowing occurred after correct responses that were misclassified as errors, indicating that conscious error awareness itself can directly trigger control adjustments. This finding has important implications for theories of adaptive control and performance monitoring. Traditionally, stronger post-error adjustments after aware errors have been interpreted as reflecting a shared underlying factor—namely, that stronger error evidence both increases the likelihood of awareness and independently drives control (Danielmeier & Ullsperger, 2011; Klein et al., 2007; Klein et al., 2013; Ullsperger et al., 2014). The present results challenge this view. We speculate that this effect might reflect a genuine action slip in the performance-classification response itself. Here, the participants mistakenly classify a correct response as an error and subsequently detect this misclassification (potentially consciously). This detection process alone could trigger the post-error slowing effect we saw in our study. Because the latency from the classification response to the next trial is shorter than from the primary task response, the resulting post-error slowing effect would appear stronger. Importantly, we are unable to determine whether participants attempted to correct such misclassifications via additional button presses, limiting our ability to directly test this account. Alternatively, the pattern could reflect a dissociation between partially independent monitoring systems. An implicit monitoring process may correctly register the primary response as accurate, while noise or instability in a higher-level evaluative system generates a conscious experience of having made an error. Such a mismatch would create a large prediction error or surprise signal at the metacognitive level, which in turn could drive stronger post-error slowing. Although more speculative, this interpretation aligns with models proposing hierarchical performance monitoring systems in which conscious error awareness and lower-level performance evaluation can transiently diverge (Alexander & Brown, 2011; Dumsky et al., 2025; Nieuwenhuis et al., 2001; Ullsperger et al., 2014; Wessel, 2012).

Moreover, we observed a post-error decrease in accuracy, which was most pronounced following unaware errors. On the one hand, this is in contrast to evidence suggesting that error commission engages both adaptive control and maladaptive orienting processes that together yield a net adaptive effect (Fischer et al., 2018)—namely, a reduced likelihood of repeating mistakes. On the other hand, this pattern aligns with recent reports demonstrating post-error performance costs rather than benefits, including evidence that errors can transiently destabilize ongoing task representations (Adkins et al., 2024). Within this framework, error awareness may serve to interrupt ongoing processing and initiate a reconfiguration of control settings. When awareness is absent, this reconfiguration may fail, resulting in continued or even worsened performance.

In summary, our data suggest that an error signal is necessary but not sufficient for behavioral adaptation. Instead, it is the **subjective interpretation** of that signal, its breaching of conscious awareness and subsequent labeling as “incorrect”, that directly gates the recruitment of control mechanisms responsible for post-error adaptations.

### Sleep deprivation may alter awareness-dependent post-error control

Sleep deprivation further altered these awareness-dependent control mechanisms following an error. While perceived errors elicited robust post-error slowing across conditions with stronger slowing for the WR condition, sleep-deprived participants showed poorer subsequent accuracy following subjectively perceived errors on objectively correct trials. Although this accuracy effect did not reach statistical robustness, it converges with recent findings suggesting that sleep deprivation compromises—rather than abolishes—post-error adjustments and may shift the balance between cautious responding and performance efficiency (Cao et al., 2025; Yue Zhang et al., 2025).

Together, these findings suggest that under sleep deprivation, awareness signals may become less reliable or less effectively integrated into cognitive control processes, leading to instable cognitive performance. As attentional resources decline, subjective evaluations of performance may increasingly misguide control processes, leading to exaggerated slowing or maladaptive changes in accuracy. Thus, sleep loss appears to compromise not only the detection of errors but also the effective use of awareness signals for behavioural regulation.

### Limitations and future directions

Several limitations should be noted. First, although the task reliably elicited aware and unaware errors, it remains a laboratory paradigm with an overall very high accuracy. Generalization to real-world settings should be approached cautiously. Second, while the present study focused on behavioural measures, future work should extend this task by adding neurophysiological recordings to directly link error awareness to error-related brain signals under sleep deprivation. Recent studies have shown that both preconscious (ERN) and conscious (Pe) error-related neural responses are reduced following sleep deprivation (Boardman et al., 2024; Boardman et al., 2021; Yue Zhang et al., 2025), offering a potential neural account for the reduced error awareness observed in our study. However, the neurocomputational mechanisms underlying diminished error awareness remain poorly understood. To further disentangle the within-trial processes that give rise to incorrect subjective performance judgments, future studies could leverage neurally informed evidence-accumulation models that jointly capture choice accuracy, response time, and subjective confidence within a single accumulation process (Duffy et al., 2025; Grogan et al., 2023; Murphy et al., 2015; Yeung & Summerfield, 2012). These models propose that both decisions and subsequent confidence judgments arise from a shared evidence accumulation process, which is reflected in the centro-parietal positivity (CPP)—a well-established neural marker that exhibits a gradual, evidence-dependent, and reaction-time–predictive build-up during decision formation (O’Connell et al., 2012; O’Connell & Kelly, 2021; Rogge et al., 2022). Using this framework, future studies could directly compare decision-formation dynamics underlying subjective performance judgments following aware versus unaware errors and test whether sleep deprivation modulates key model parameters and their neural proxies, such as decision boundaries or CPP amplitude around the time of self-evaluation. Third, the present study examined total sleep deprivation; whether similar mechanisms operate under chronic sleep restriction remains an important open question. Notably, recent evidence indicates that sleep restriction can produce stronger negative effects on behavioural error awareness and neural markers of performance monitoring than a single night of total sleep deprivation (Boardman et al., 2024).

## Conclusion

In summary, one night of total sleep deprivation reduces behavioural error awareness and disrupts post-error adjustments in a time-dependent manner. Most importantly, our findings demonstrate that adaptive control is strongly shaped by subjective error awareness, even when that awareness is inaccurate. These results highlight conscious performance evaluation as a key mechanism linking sustained attention, sleep loss, and cognitive control, underscoring the importance of considering subjective awareness when studying adaptive behaviour under fatigue.

## Acknowledgments

We thank all our participants who took part in this research for the generosity of their time and commitment. We also want to acknowledge the hospitality from the Leibniz Institute for Neurobiology in Magdeburg that allowed us to conduct our study there. And we thank Luca Budinger for her extraordinary support during data collection. This research was supported by the Deutsche Forschungsgemeinschaft, Grant/Award Number: SFB 1436, sub-project C04 (to JT and MU) and the European Research Council, Grant/Award Number: 101018805 (to MU).

## Supplementary Results

**Supplementary Figure 1.**
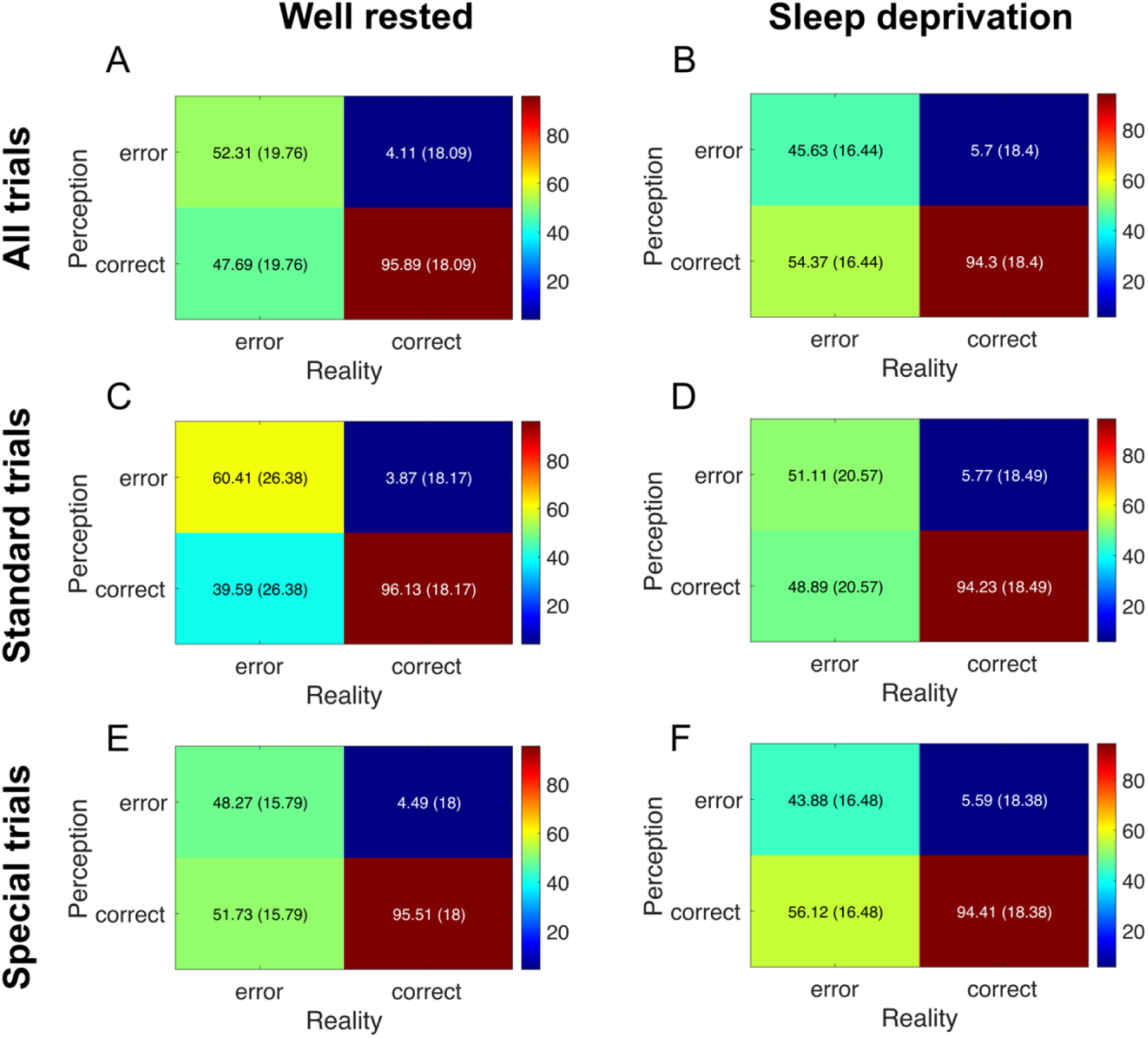
Self-evaluation confusion matrix. Here we show the percentages of responses according to signal-detection theory to illustrate participants’ ability to discriminate between correct and error responses across all trial types **(A-B)**, standard trials **(C-D)**, and special trials **(E-F)**. Overall, participants demonstrated good discrimination between actual correct and error responses. Yet, a substantial proportion of actual error responses remained unnoticed. This pattern of results suggests that a) participants were able to successfully evaluate their responses and that b) the task induced a sufficient number of unaware errors, enabling us to study the effect of error awareness on cognitive control processes and how this relationship might be affected by sleep deprivation. Note that sleep deprivation appears to reduce the error detection sensitivity particularly on standard trials **(C&D)**.

### Time on task effects analyses

To examine time-on-task effects, we first computed a smoothed density estimate of key task performance variables (error rate, error awareness rate, and VCT) using a Gaussian kernel with a full width at half maximum (FWHM) of 9 trials. This procedure effectively integrated information from the surrounding 20 trials via a weighted average of the performance measures. Next, we discretized the available trials into 100 quantiles. Finally, we plotted the difference (Δ = WR − TSD) to visualize how sleep deprivation affected each performance variable over the course of the task.

To formally test the effect of sleep deprivation on these variables, we applied a sliding-window approach (step size = one quantile; window size = 25 quantiles), following recent recommendations by Jenkins and Quintana-Ascencio (2020). For each window, we fitted a separate hierarchical model to estimate the effect of sleep deprivation on the respective performance measure. The basic model was defined as follows:

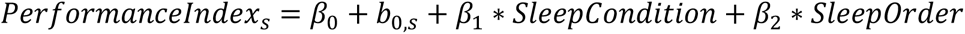

where *βs* are fixed effects, *bs* denotes the subject-specific random intercepts, SleepCondition indicates the state in which the task was performed (coded as −1 for well-rested and 1 for sleep-deprived), and SleepOrder is a control regressor accounting for session order (sleep deprivation in the first vs. second session, coded as −1 and 1, respectively). The results of these analyses are shown in Supplementary Figure 2:

**Supplementary Figure 2.**
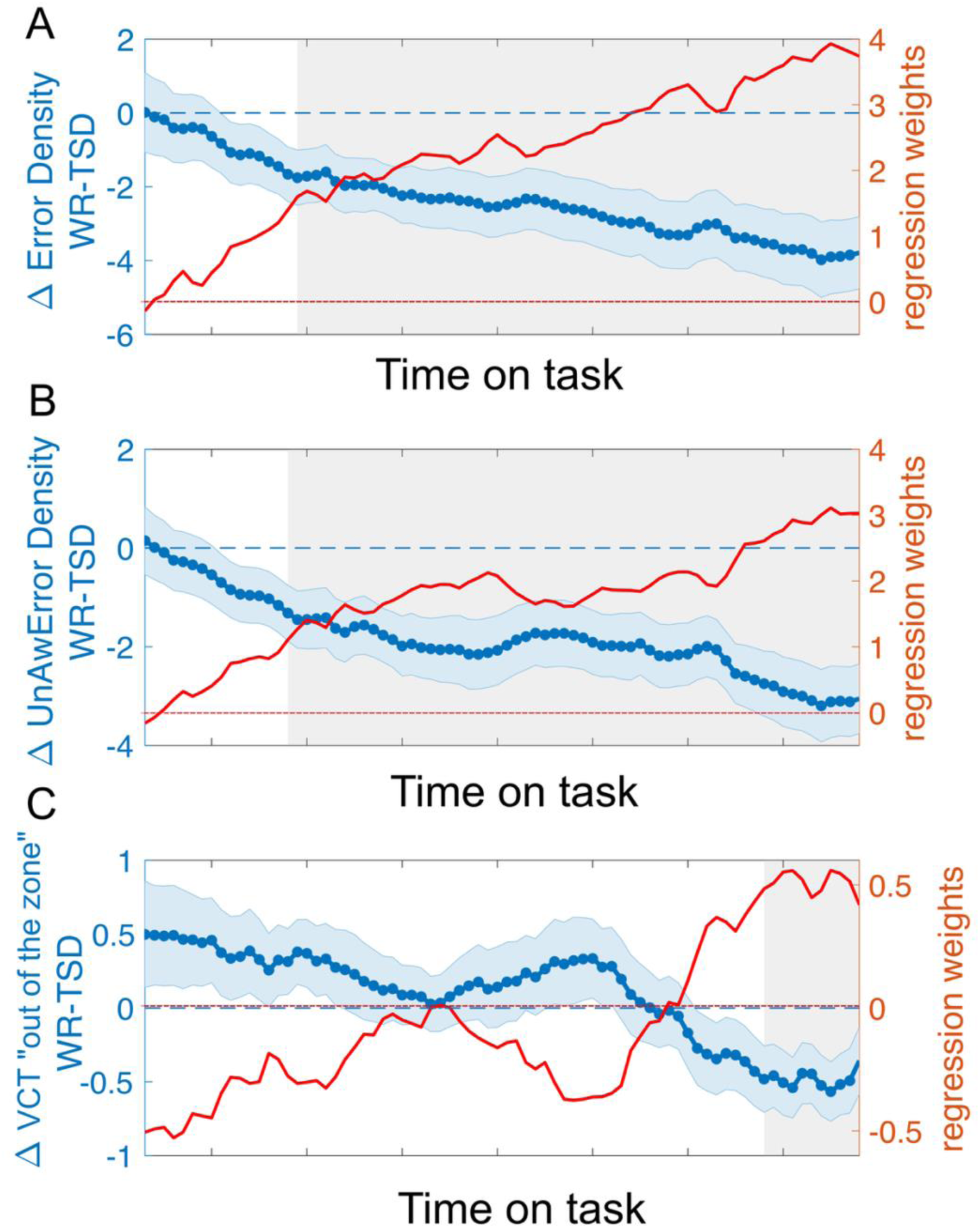
The effect of sleep-deprivation depends on time on task. There were no performance differences between sleep deprivation and rested wakefulness at the beginning of the task but during the course of the task, total error **(A)** and unaware error densities **(B)** increased significantly in the sleep deprivation condition. Similarly, at the end of the task, sleep deprivation increased the time spent in the out of the zone state **(C)**. Gray areas indicate time points where there was a meaningful effect of sleep deprivation (i.e., the credible interval excluded zero). Beta values of that model (WR – STD) are depicted in red. *Note:* Gray areas indicate time points at which there was a meaningful effect of sleep deprivation (i.e., the credible interval excluded zero). Beta values from the model (WR – SD) are depicted in red. The red dashed line indicates a beta value of zero; the blue dashed line indicates zero difference in behavioural indices. Blue shaded areas represent the SEM.

